# Bivariate spatial point patterns in the retina: a reproducible review

**DOI:** 10.1101/029348

**Authors:** Stephen J. Eglen

**Affiliations:** Cambridge Computational Biology Institute, Department of Applied Mathematics and Theoretical Physics, University of Cambridge, CB3 OWA

## Abstract

In this article I present a reproducible review of recent research to investigate the spatial positioning of neurons in the nervous system. In particular, I focus on the relative spatial positioning of pairs of cell types within the retina. I examine three different cases by which two types of neurons might be arranged relative to each other. (1) Cells of different type might be effectively independent of each other. (2) Cells of one type are randomly assigned one of two labels to create two related populations. (3) Interactions between cells of different type generate functional dependencies. I show briefly how spatial statistic techniques can be applied to investigate the nature of spatial interactions between two cell types. Finally, I have termed this article a ‘reproducible review’ because all the data and computer code are integrated into the manuscript so that others can repeat the analysis presented here. I close the review with a discussion of this concept.

---

**Table of abbreviations**
Term: meaning
BC: blue cone
BCBP: blue cone photoreceptor
CSR: complete spatial randomness
GCL: ganglion cell layer
INL: inner nuclear layer
IPL: inner plexiform layer
ONL: outer nuclear layer
OPL: outer plexiform layer
RGC: retinal ganglion cell

## Introduction

The retina is the neural structure located at the back of the eye. Light passes through the eye and is converted into electrical activity by the photoreceptors. The retina then performs sophisticated processing of the visual scene before transmitting this information to the brain for further processing. See Wässle (2004) for an overview of this processing. Estimates vary, but there are thought to be at least 60 types of retinal neuron (Masland, 2012) that perform unique computations to transform the visual scene into a neural code. The computations performed by each neuron depend critically upon who it receives inputs from; to a first approximation, retinal neurons communicate with other neurons that are close by in adjacent neuronal layers. The spacing of retinal neurons therefore influences the computations performed within the retina. Furthermore, retinal neurons of a given type are usually distributed across the retina to ensure that all regions of the visual scene can be efficiently processed.

The cell body of each type of neuron typically is located in just one of the three nuclear layers (Figure 1). Although the cell body of a retinal neuron has a three dimensional position (*x*, *y, z*) we can often disregard *z* for two reasons. First, once we know the type of a neuron, its *z* value is fairly well determined. Second, the range of *z* (given by retinal thickness, about 0.2 mm; Ferguson et al., 2013) is much smaller than the range of *x* and *y* (4-5 mm Sterratt et al., 2013) Therefore in this review, we will treat neuron positions as two-dimensional.

**Figure 1:**
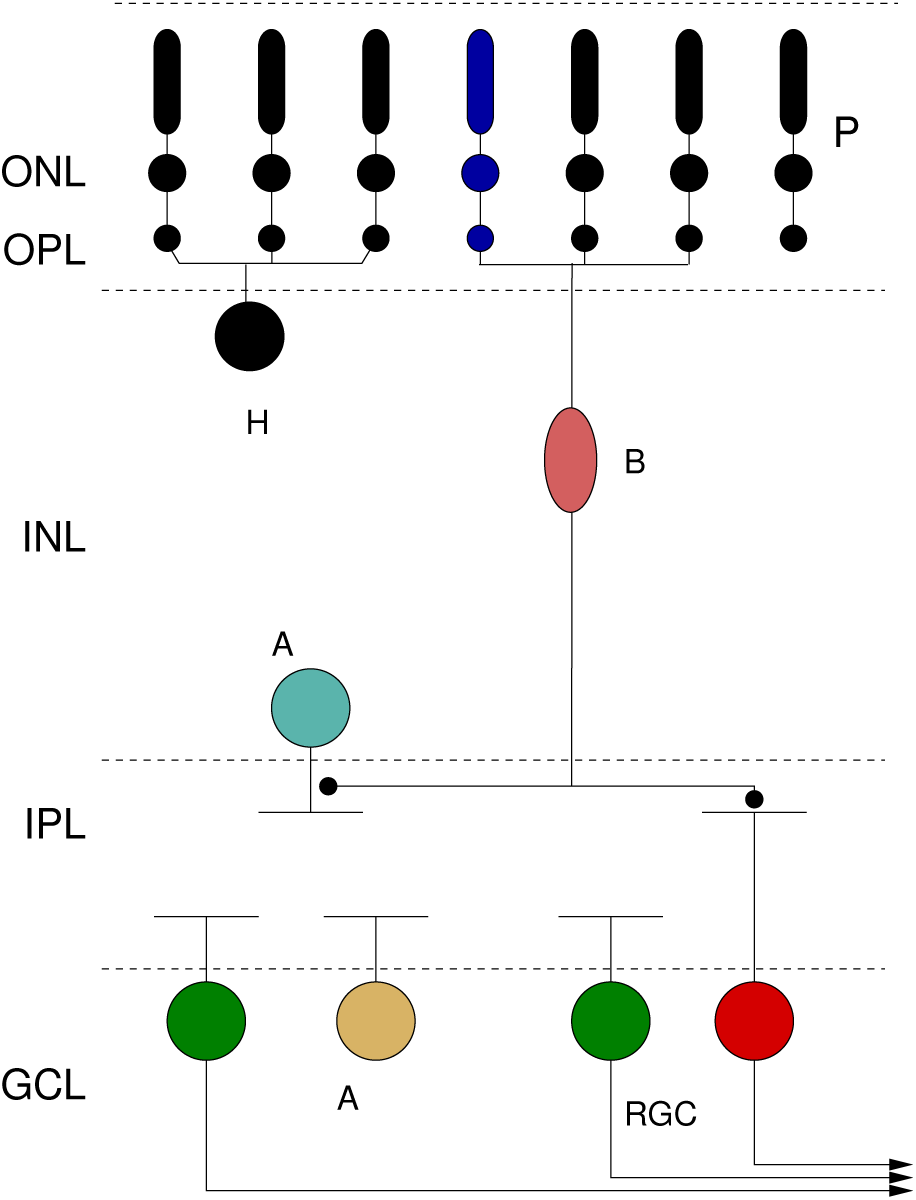
Cross section of vertebrate retina. The cell bodies of neurons are located in three layers: outer nuclear layer (ONL), inner nuclear layer (INL) and ganglion cell layer (GCL). Connections between these neurons are typically formed in the outer plexiform layer (OPL) and inner plexiform layer (IPL) that separate the three layers. Neurons of a given type normally occupy just one layer. The diagram indicates the five main classes of neuron: photoreceptor (P), horizontal cell (H), bipolar cell (B), amacrine cell (A) and retinal ganglion cell (RGC). If a slice is taken through one layer, e.g. GCL, we can observe the spatial regularity of neurons within one layer (Figure 2). This figure adapted from Figure 1 of Eglen (2012).

One organisational feature of the retina found in many species is that of “retinal mosaics”: neurons are typically positioned within a layer in a semi-regular manner. Figure 2A shows a typical example from cat retina. Figure 2B shows the result of a corresponding model, which will be explained below. These patterns visually appear to have similar spatial properties. The spatial organisation of point processes such as these can be quantified using the *K* function (Ripley, 1976), which calculates the expected number of points within a given distance of any point. The *K* function can analyse both univariate patterns (cells of one type; Figure 2) or bivariate patterns (cells of two types; Figure 4). Given that the univariate *K* function is a simplification of the bivariate *K* function, we first define the bivariate function.

**Figure 2:**
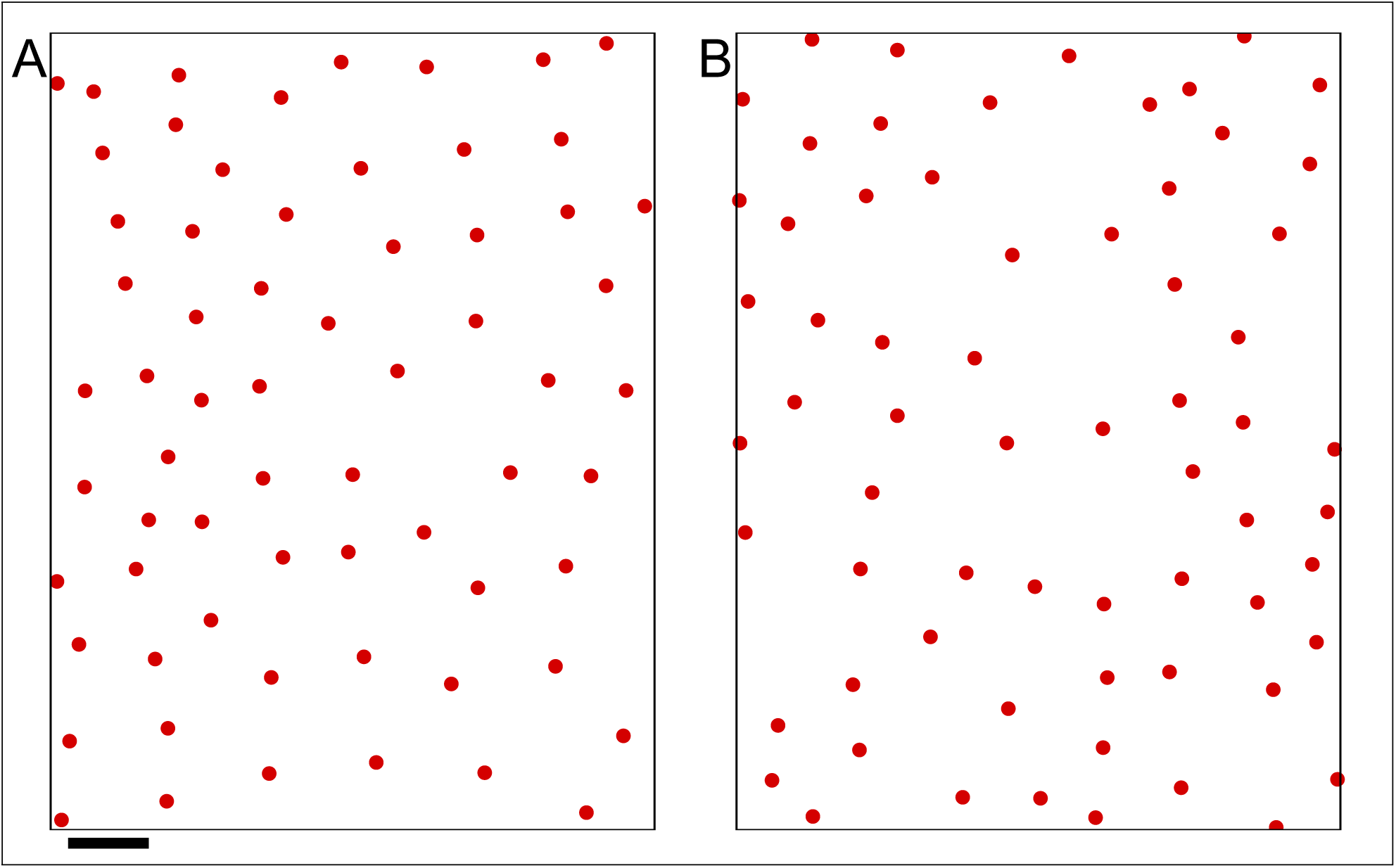
An example retinal mosaic: beta on-centre retinal ganglion cells (Wässle et al., 1981). On the left is the observed map, and the right is an example univariate simulation with matching field and density of points. Scale bar is 100 μm; soma are drawn to scale with a radius of 9 μm.

We store the position of the *i*th type 1 neuron as a two-dimensional vector x_1_*_i_* where *i* ∈ [1, *n*_1_] and likewise for the *j*th type 2 neuron in x_2_*_j_*, *j* ∈ [1, *n*_2_]. We then define the bivariate *K* and *L* functions as

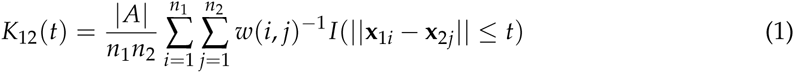

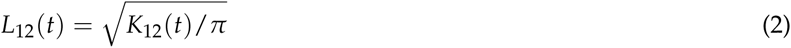

The weighting factor *w*(*i, j*) is a normalisation term to account for edge effects. It is defined as the fraction of circumference of the circle centred at x_1_*_i_* and with radius ||x_1_*_i_* − x_2_*_j_* || that lies within the sampling window *A. |A*| denotes the area of the sample window. The univariate *K* function (to analyse one set of points) is calculated by defining *n*_2_ = *n*_1_ and x_2_*_i_* = x_1_*_i_* for all points and ignoring self-counts.

The *L* function is a rescaling of the *K* function. For either the univariate or bivariate L function, if for a range of distances *t* we find that *L*(*t*) ≈ *t*, this indicates that there is no spatial order in the point pattern. This is commonly termed *Complete Spatial Randomness* (CSR) (Diggle, 2002). If however *L*(*t*) > *t*, this would indicate *spatial clustering:* points are positioned close to each other more often than chance. Finally, *L*(*t*) < *t* implies *spatial regularity,* whereby points avoid each other (Diggle, 2002).

Many other methods for quantifying order in retinal mosaics have been proposed, ranging from simple nearest-neighbour methods (Wässle and Riemann, 1978) to those based on Voronoi diagrams (Keeley and Reese, 2014). Several of these measures have been empirically compared (Cook, 1996). In this review we focus on probably the single most useful measure, the *K* function. It is worth noting that a closely related variant of the *K* function, the density recovery profile, was proposed specifically for the analysis of retinal mosaics (Rodieck, 1991). However, as noted by its author, the density recovery profile is effectively the derivative of the *K* function computed with a given bin width, and has the disadvantage that the bin width must be specified.

As can be seen in Figure 3, retinal mosaics normally show evidence of spatial regularity over small distances, in this case up to around 150 μm. Beyond this distance, there is no further order and so the *L* function matches that expected from CSR. Hence, this suggests there is a local exclusion zone working over relatively small distances (no more than 150 μm here) to space neurons apart. Several biological mechanisms have been proposed to account for the formation of such exclusion zones, including lateral inhibition of cell fate (Takesue et al., 1998), lateral migration of neurons (Reese and Galli-Resta, 2002) and cell death (Jeyarasasingam et al., 1998; Resta et al., 2005).

To check that the global patterns of retinal mosaics can emerge from simple local exclusion zones, we can build models whereby cells are positioned subject to such local rules only. The simplest of these models is called the *exclusion zone* model, or the d_min_ model (Eglen, 2012). In this model, cells are added one-by-one into the array at random locations as long as they are some distance d_min_ away from all previously positioned cells. The larger the value of d_min_ the bigger the exclusion zone surrounding each cell. d_min_ is the only free parameter of the model, and is often varied from cell to cell by sampling values from a Normal distribution. In this model, we treat the neuron as simply a point in space which represents the centre of the soma. Although adult neurons have extensive dendritic and axonal arborisations, our simplification of regarding the neuron as a point is appropriate as the cellular spacing occurs early enough in development, when axons and dendrites are relatively sparse, if present at all (Reese et al., 1999).

To test whether the d_min_ model accounts for the data, we need a method to compare model output with experimental data. One common approach used in spatial statistics is to compare their *K* or *L* functions either for one run, or many runs, of the model (Figure 3). Informally, if the *L* function for the data falls within the envelope of the *L* functions from multiple runs of the model from different initial conditions, then we say that the model fits the data. More formally, empirical p values can be computed to compare model and data (Diggle, 1986). In Figure 3 there is a good qualitative match given that the data fall mostly within the bounds of the confidence intervals. There is a minor excursion of the data outside the confidence intervals around 130–150 μm however which suggests the fit is not perfect. Further tuning of the parameters, or switching to an alternative model with combined hard-core and soft-core exclusion zones (Diggle, 2002), might improve the fit slightly.

Given that the d_min_ model is a phenomenological model, it suggests only that local exclusion zones are sufficient to recreate global order, but cannot tell us which developmental mechanisms are responsible. To assess which mechanisms might be responsible, we need to examine mechanistic models (Eglen et al., 2000; Eglen and Willshaw, 2002; Ruggiero et al., 2007). However, although we cannot use the current methods to understand the mechanisms of development, the existence of retinal mosaics is often used as an indicator of an independent cell type (Cook, 2003; Seung and Sümbül, 2014). Studying the nature of the retinal mosaics is therefore also helpful to understanding how to classify neuronal types.

**Figure 3:**
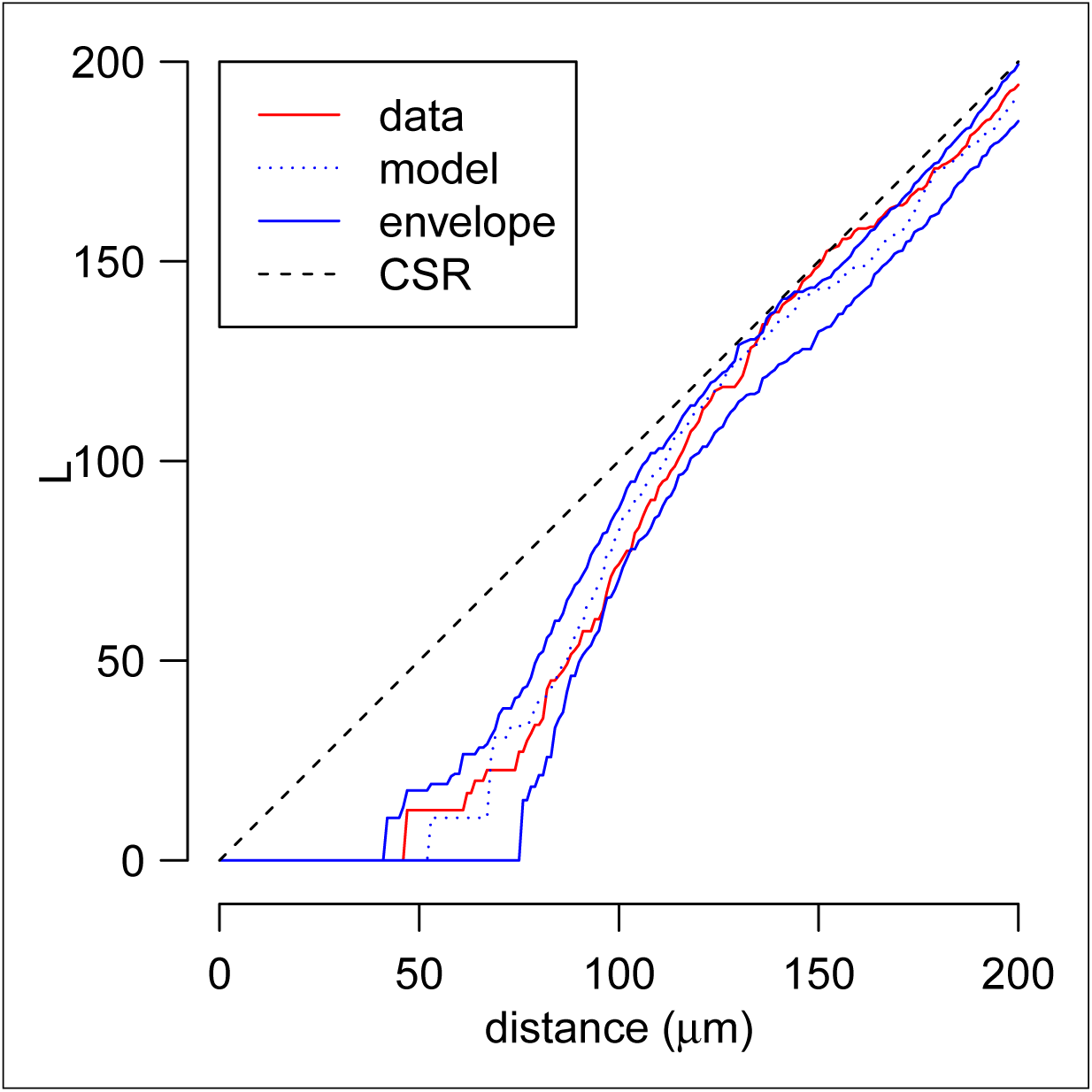
Evidence for local exclusion zones in retinal mosaics. The solid red curve shows the *L* function for the mosaic shown in Figure 2A. The dotted blue curve is the corresponding *L* function for the model output in Figure 2B; the solid blue curves are the confidence intervals from the model estimated by finding the range of *L* functions from 99 simulations with fixed parameters but different initial conditions. The dotted black line is the expectation for the *L* function assuming complete spatial randomness (CSR).

## Bivariate patterns

In Figures 2 and 3 we have explored the spatial relationships among cells of the same type. We now consider the spatial relationships between cells of two different types. In particular, we examine several examples of related pairs of neuronal types. Perhaps the most famous example of such a bivariate pattern is the on-centre and off-centre beta retinal ganglion cells (Wässle et al., 1981). These two types of neuron differ physiologically in terms of whether they respond to increases or decreases in light levels in the centre of their receptive fields. Figure 4A shows an example of a bivariate point pattern from cat retina, and Figure 4B shows a stochastic model again based on exclusion zones (to be described later). How do we decide what structure might be present in the data, and whether the model is a good fit for the data? Diggle (1986) established a quantitative framework to examine these questions nearly thirty years ago. In this review we apply this framework to datasets from three pairs of retinal neuronal types that suggests different pairs of neuronal types are organised in different ways. This suggests that the developmental mechanisms generating these patterns are not universal, but depend on which pairs of cells are being considered. Functional independence is the null hypothesis that says there are no developmental interactions between pairs cell types, whereas random labelling and functional dependence are alternative hypotheses suggesting that developmental mechanisms drive their bivariate patterning. In this review we use the phrase coined in Diggle (1986) of “bivariate spatial point patterns” to refer to the spatial arrangement of two types of retinal neuron within the same sample window.

**Figure 4:**
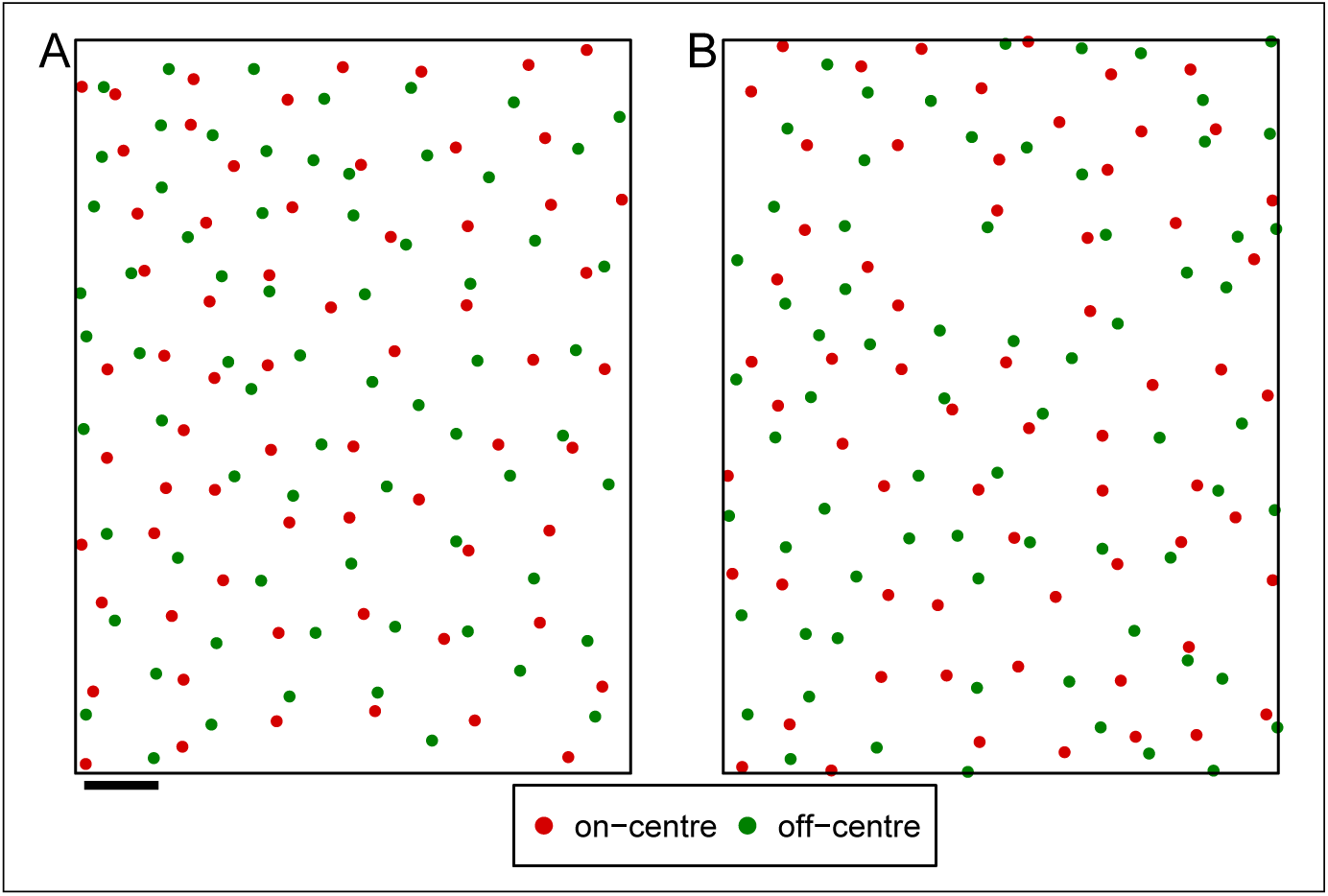
Bivariate spatial point pattern. A: observed field of cat on- and off-centre beta retinal ganglion cells, W81S1, from (Wässle et al., 1981); B: simulation using bivariate PIPP rule. Scale bar is 100 μm; soma are drawn to scale with a radius of 9 μm.

## Example one: evidence for functional independence

With multiple types of neurons present in the retina, it is possible that neurons of different types constrain each other in their spatial positioning. Interactions between two types of neurons are termed *heterotypic interactions*, to be compared with interactions between neurons of the same type, called *homotypic interactions*. Discovering the interactions present among cell types can help understand the developmental mechanisms that create neural circuits.

The simplest situation is that of *statistical independence* between two spatial point patterns, which means that the spatial location of one type of neurons is completely unaffected by the spatial location of another type of neurons. Statistical independence between type of neurons occupying different layers of the retina was originally reported for cholinergic amacrine cells (Diggle, 1986) and subsequently for several other pairs of neuron in rabbits (Rockhill et al., 2000) and zebrafish (Mack, 2007). This suggests that homotypic interactions are sufficient to create the spatial organisation within each type of neuron.

Full statistical independence between pairs of neurons can be shown only when the two types of neurons occupy different layers of the retina. When both types of neurons are in the same layer, their physical size constraints need to be taken into account as neurons cannot occupy the same physical space. The cell bodies of retinal neurons are typically at least 10 μm in diameter for example, which introduce steric hindrance effects (Cook, 2003) between all neurons, irrespective of their type. For example, cell bodies of many types of retinal ganglion cell (RGCs) are located in the ganglion cell layer (GCL). Given that there are at least fifteen anatomical types of RGC in mouse retina (Sümbül et al., 2014) these steric hindrance effects are likely to be significant.

To account for these physical effects, we devised the term *functional independence* to describe the case when two spatial point patterns do not interact, except for these short-range steric hindrance effects (Eglen et al., 2006). To test for functional independence, we examined in detail the on- and off-centre beta RGCs in cat (Figure 4; Eglen et al., 2006). We extended the exclusion zone model presented earlier such that were effectively three d_min_ exclusion zones in the model, termed a pairwise interaction point process model (Ripley, 1976, 1977). The first exclusion zone, 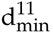, was for modelling homotypic interactions between pairs of on-centre neurons, and likewise a second exclusion zone, 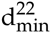, for homotypic interactions between off-centre neurons. The third d_min_ exclusion zone, 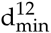, was used for modelling the heterotypic interactions between an on-centre and an off-centre neuron. The effective values of 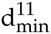 and 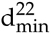 for homotypic interactions was quite large, around 50-100 μm. By contrast, we set 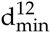 to the relatively small value of 18 μm — the average diameter of RGC soma. An example of the model output using this bivariate model is shown in Figure 4B. This model was judged to be a good fit to the data when comparing various spatial statistics of the observed data and model simulations (Eglen et al., 2006). We therefore concluded that the spatial positioning of on- and off-centre beta RGCs are functionally independent of each other.

## Example two: evidence for random labelling

The framework by Diggle (1986) allowed the *random labelling* hypothesis to be tested. The random labelling hypothesis is simply that one homogeneous population of cells is created with one exclusion zone constraint. This population then divides into two, without cells moving again, by simple random labelling: each cell is independently labelled as type 1 or type 2 with a fixed probability. There is therefore only one exclusion zone constraint in random labelling, compared to two (usually different) exclusion zones constraints when creating cells using functional independence. The random labelling hypothesis was originally rejected for the cholinergic amacrine cells (Diggle, 1986; Diggle et al., 2006). However, in recent years, data consistent with random labelling has appeared in both ferret retina (Eglen et al., 2003) and in Drosophila (Bell et al., 2007).

For example, in the ferret retina, dopaminergic amacrine neurons are found in two separate layers of the retina, one set in the inner nuclear layer (INL) and the other in the GCL (Eglen et al., 2003). It was unclear whether these cells in the two layers form two distinct types, or belong to one type of neuron. Figure 5 demonstrates the random labelling hypothesis, showing one real field, along with five random realisations where for each cell, its location is preserved, but the type of the cell (GCL versus INL) is randomly switched, subject to preserving the overall ratio of cells in each sub-type.

By visual inspection alone, it would appear that all six fields appear similar, which informally suggests that the data may be consistent with random labelling. (The experimental data are shown as example five in Figure 5; the remaining examples in Figure 5 are five examples of random relabelling of the data.) This is confirmed by calculating the *L*_12_ function for the experimental data and 99 realisations of random labelling. The *L*_12_ function for the data mostly remains within the envelope formed by the 99 simulations (Figure 6). (There is a modest departure of the data from the envelope at about 100–150 μm which might indicate the parameters could be further tuned to improve the fit.) From this, we can conclude that although the cell bodies of dopaminergic amacrine cells occupy two distinct layers, both layers of cells should be considered together as a single type of neuron.

**Figure 5:**
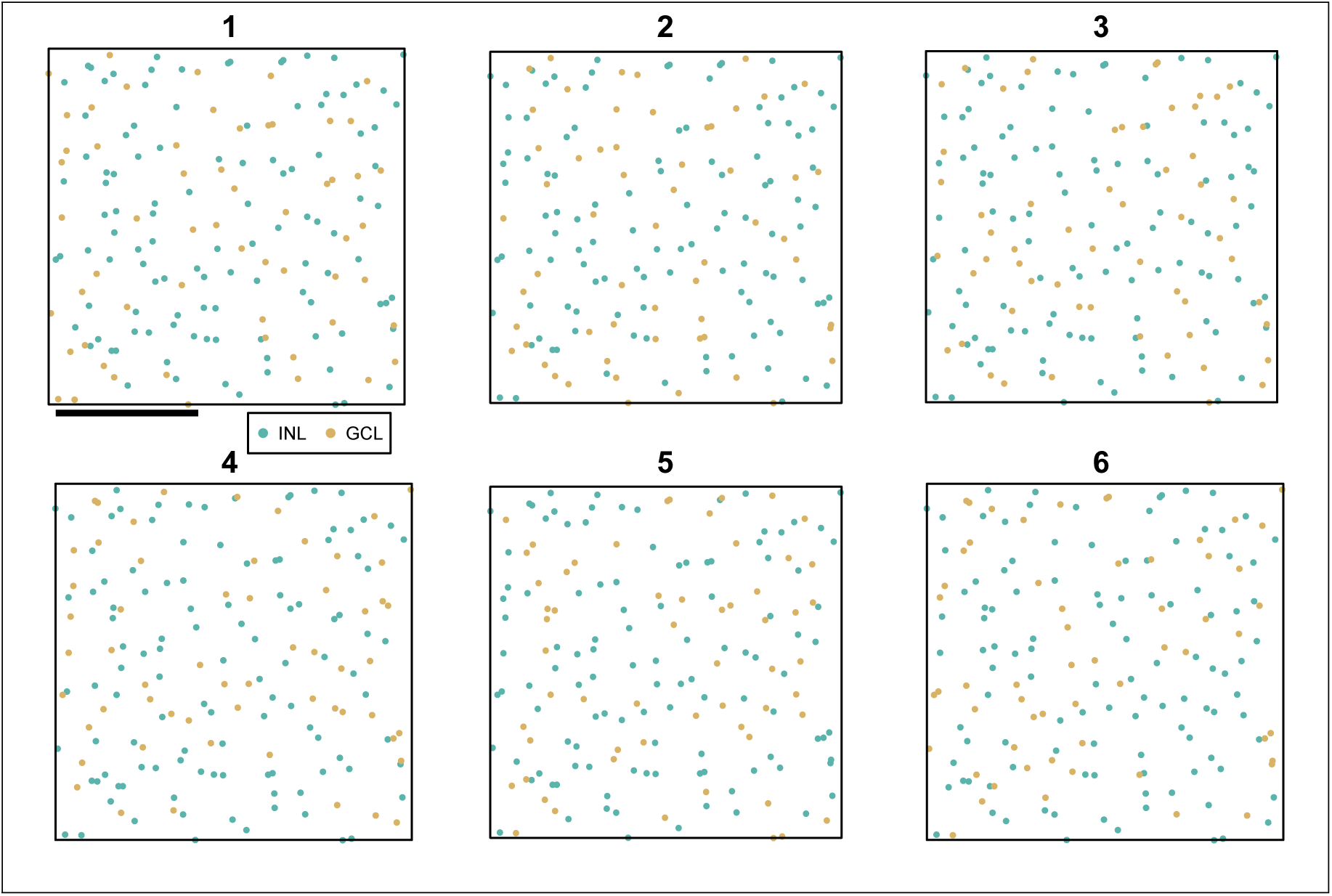
Dopaminergic amacrine neurons in ferret, showing one real field along with five simulated fields. Which figure is the experimental data? Scale bar is 1 mm; soma are drawn to scale with a radius of 22 μm.

**Figure 6:**
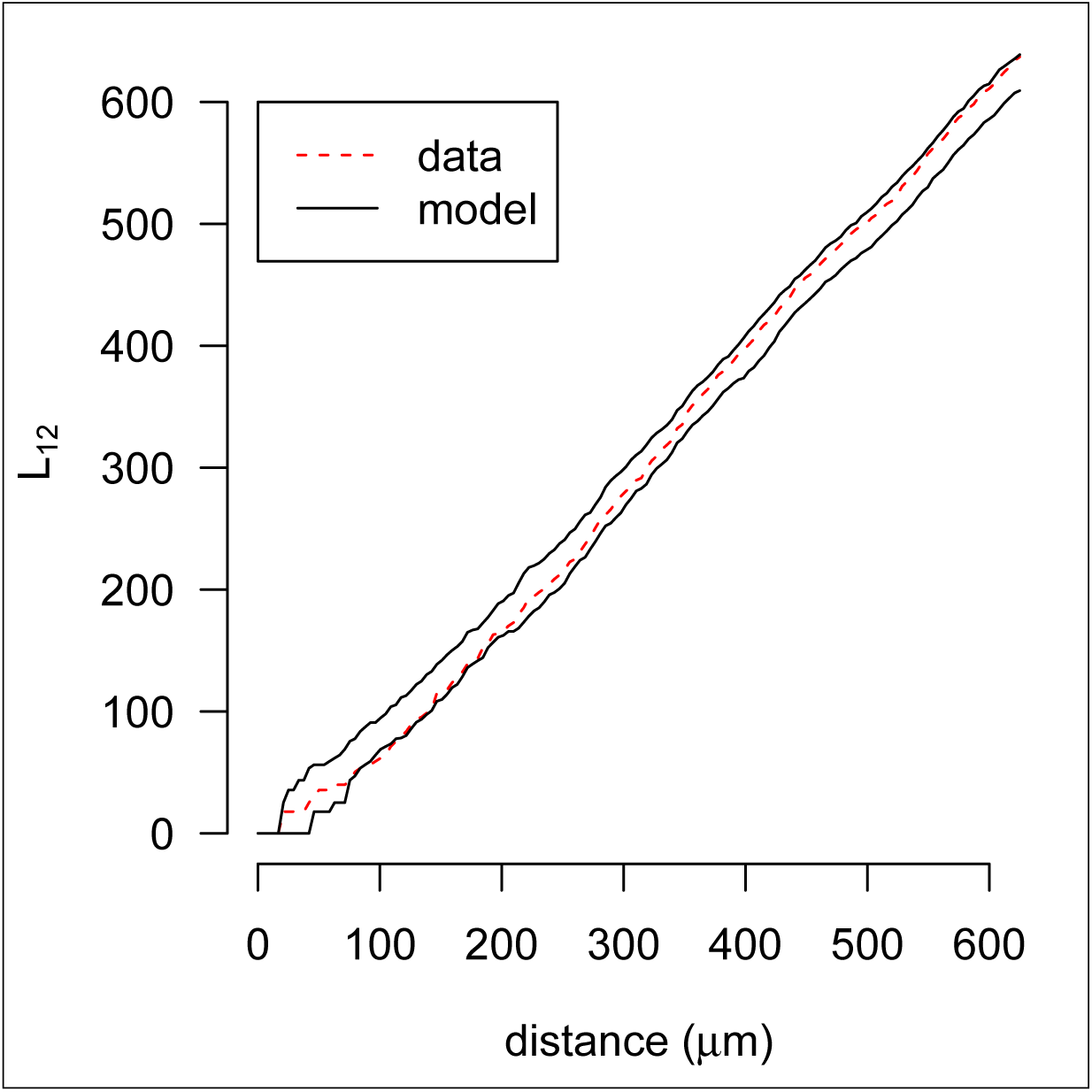
*L*_12_ function for dopaminergic amacrine data set shown in Figure 5. The *L*_12_ function for the observed data (example 5 in Figure 5) is shown as a dotted red line; black lines show the envelope from 99 simulations of random relabelling.

## Example three: evidence for functional dependence

Many researchers have tested for the existence of functional dependencies between two types of retinal neuron. In particular, a systematic study examining ten pairs of cross-correlations from five types of retinal neuron failed to reveal any cross-correlation (Rockhill et al., 2000). The most prominent example of cross-correlation to date has been found in macaque, comparing the spatial location of blue cone photoreceptors (BC or “blue cones”) and their synaptic targets in an adjacent layer, the blue cone bipolar cells (BCBP) (Kouyama and Marshak, 1992, 1997). An example of such a mosaic is shown in Figure 7. These cells tend to cluster together, as shown by the increased values of the *L*_12_ function over distances up to around 25 μm. As BCBPs receive input only from BCs, it would make sense for them to be close to each other, to minimise the synaptic wiring required. Our preliminary modelling studies suggest that we can account for these spatial patterns by fixing the locations of blue cones, and allowing the BCBPs to make contact with nearby blue cones and then move to reduce tensile forces within dendrites (Lønborg, 2008), but other mechanisms might also be involved.

This example of BC and BCBPs demonstrates a spatial clustering between two types of neuron. By contrast, there has been one report of a negative correlation (spatial avoidance) between two types of neuron in three related species from the cat family (Ahnelt et al., 2000). In this example, one class of axonless horizontal cells are found at significantly lower densities in locations near to blue cones. Unfortunately however, we do not have the data from this study to investigate directly the nature of the *L*_12_ function.

**Figure 7:**
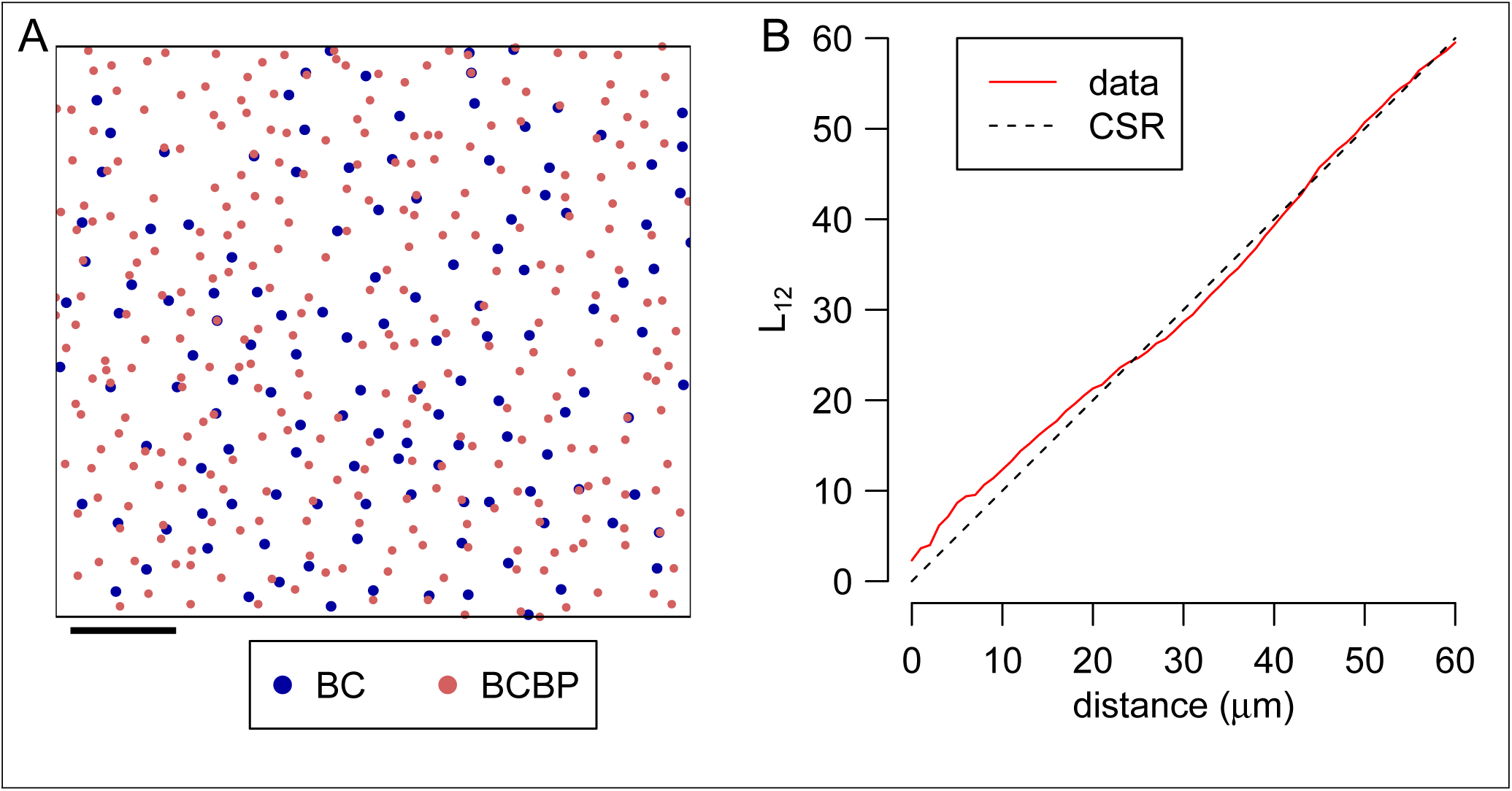
Functional dependence between two types of neuron. A: An example field of blue cones (BC; blue) and blue cone bipolar cells (BCBP; red). This is field one from the dataset provided by Kouyama and Marshak (1997). Scale bar is 100 μm; soma are drawn to scale with a radius of 5 μm (BC) and 4 μm (BCBP). B: The L_12_ function for the data shown in A (solid red line) and expectation for the curve under Complete Spatial Randomness (CSR).

## Future directions

That these two examples of positive and negative cross-correlation both involve blue cones may be a coincidence or may reflect something about the developmental mechanisms that constrain the movement and positioning of neurons. In either case, these two examples are clear examples of where further modelling and analysis might help our understanding of the development of the retina. Furthermore, we anticipate that in the near future, as more genetic markers for unique cell types in mouse are carefully characterized (Sümbül et al., 2014), I expect that there will be many more opportunities for labelling multiple types of neuron within individual samples. The simultaneous localisation of five types of cone photoreceptor within a field has already been achieved in chick retina (Figure 8; Kram et al., 2010). In this case, the identity of cone photoreceptors was revealed by examining the type of oil droplet found just beneath their outer segment. By examining all pairs of neurons, no evidence for spatial correlation was found (Kram et al., 2010). However, this does not rule out that there might be higher-order dependencies within the multiple cell types.

**Figure 8:**
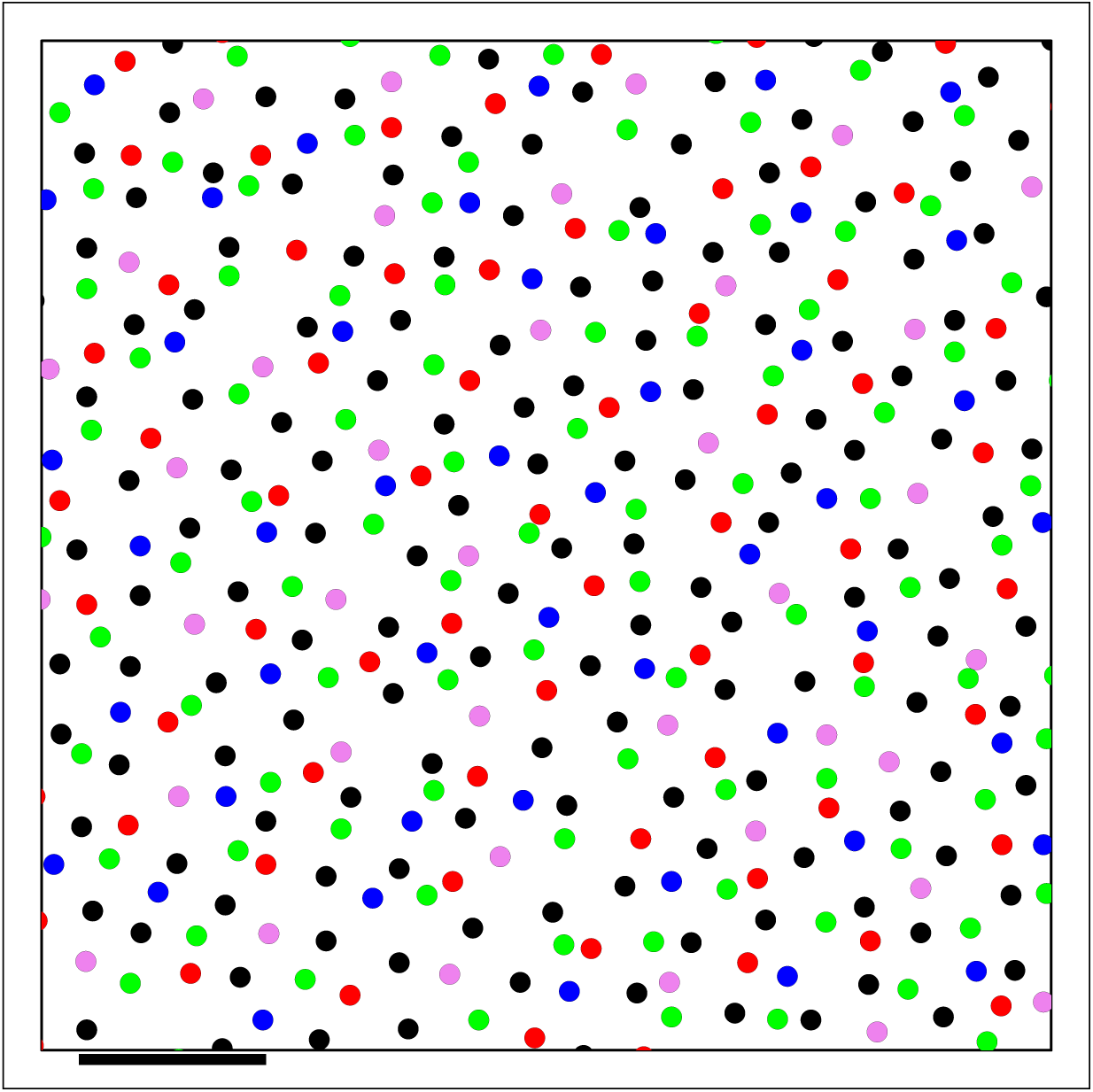
The spatial layout of five types of cone photoreceptor from chick retina (Kram et al., 2010). The colour of each neuron represents the type of neuron (red, green, blue, violet) except double-cones are coloured black here. Scale bar is 20 μm; soma are drawn to scale with an estimated radius of 1.1 μm. Data shown here is “DN field 4” taken from Supplementary Information of Kram et al. (2010).

The retina has been chosen here as a model system for exploring spatial correlations in the nervous system. The retina is particularly useful for this task because of its strongly laminar structure and the availability of markers for selectively identifying cells of one type. However, we imagine these techniques will have utility in other regions of the nervous system, and have indeed been applied to prefrontal cortex, striatum and spinal cord (Cotter et al., 2002; Gangarossa et al., 2013; Prodanov et al., 2007). Finally, although we have used these techniques here to demonstrate the spatial organisation of anatomical features, they can also be used for physiological features, such as the arrangement of centres of receptive fields (Hore et al., 2012).

## Reproducible research

**Table 1:**
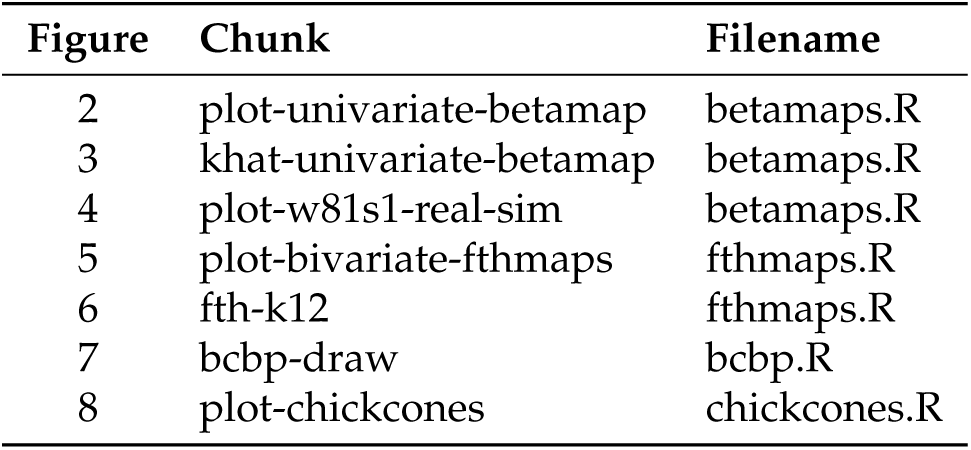
Code chunks used to generate each figure

| Figure | Chunk | Filename |
| --- | --- | --- |
| 2 | plot-univariate-betamap | betamaps.R |
| 3 | khat-univariate-betamap | betamaps.R |
| 4 | plot-w81s1-real-sim | betamaps.R |
| 5 | plot-bivariate-fthmaps | fthmaps.R |
| 6 | fth-k12 | fthmaps.R |
| 7 | bcbp-draw | bcbp.R |
| 8 | plot-chickcones | chickcones.R |

When writing a review article such as this, it is typical to include results, usually as figures or tables, that have been published before in other articles. For computational work, this typically means include the results of previous modelling or analysis. I have deliberately taken a different approach in this article: all the results presented here have been generated afresh, albeit based on my previous work, and new figures generated for the results. The one exception to this is Figure 1, which is a schematic of retinal layers. Furthermore, all the code and data required for others to generate new figures are available as part of an R package. This package is publicly available at http://github.com/sje30/eglen2015; see that URL for full details of how to regenerate this article. Table 1 describes the chunk and filename used to generate the reproducible figures in this article.

This paper is therefore the main output of the R package that I have created. This approach to writing papers is known as reproducible research, since the aim is that others should be able to reproduce the results presented in this paper for themselves. (Given that the simulations in this paper are stochastic, there will be some variation each time the paper is created.) This notion is fairly new to computational neuroscience, with Delescluse et al. (2011) possibly as the first example. However, in other disciplines, such as computational biology, reproducible research papers have a slightly longer history (Gentleman et al., 2004).

There are now several alternative workflows for generating such reproducible research documents. I have chosen to use the knitr literate programming framework for R (Xie, 2013). This provides a powerful framework for mixing documentation (the paper) and code (the analysis). Although knitr can work with several programming languages, it has the strongest support for the R programming language, which was described as the most popular tool in statistics in 2014 (Tippmann, 2015). The packaging system in R blends easily with the reproducible research document since it allows an author to include all data and code relating to a project into one package. This package can then be shared easily with other R users.

Although this packaging system can work well in R, and has done for this modest-sized review, there are still technical challenges to overcome, mostly when dealing with large data sets or with computations that take a long time to complete. Another technical problem is how to ensure that another user has a compatible environment for running your software. Running the same code in different environments may well produce different results or even fail to run. An early attempt to solve this problem was the use of virtual machines to package all the data and code relating to the project such that others could re-run the analysis (ENCODE Project Consortium, 2012). More recently, the Docker system provides a convenient way for the entire computing environment relating to a project to be efficiently captured and then run. Docker runs on all major operating systems and provides lightweight linux virtual machines and hence a convenient way of getting software to run on different platforms. For example, once Docker is installed, the entire environment required to generate this paper can be downloaded and initialised using the command:

~~~
docker run -d -p 8787:8787 sje30/eglen2015
~~~

Once this is complete, which typically takes a few minutes the first time it is run, the user can open a web browser to run R locally and then view the PDF or edit and regenerate this article. Given that the R environment is open source, the support for R in docker is fairly mature (Boettiger, 2014). Further details on running the docker system with R are given at https://github.com/rocker-org/rocker/wiki/Using-the-RStudio-image. The biggest restriction to Docker for reproducible research is that the required software must be open source (which excludes environments such as Matlab which are popular in Neuroscience). An additional concern is that both Github and Docker are private services, and public infrastructure might be preferred for long-term availability of valuable scientific resources.

Although the concept of reproducible research documents is fairly new in most mainstream journals, I hope we will see more people choosing to write their papers in this fashion in years to come. Just as in recent years journals and funding agencies have embraced the notion of “open data” so that data relating to a paper should be freely shared, there is now a growing trend for journals to ask for other research artefacts, notably computer code, to be shared. The journal *Biostatistics* was one of the first to encourage researchers to share code and data, along with reproducible research, by including suitable kitemarks (“badges”) on the front page of the article (Peng, 2009, 2011). The journal *Science* requires authors to generated supporting online material including relevant code and data. As an example, Vogels et al. (2011) provides a comprehensive supplement in the field of computational neuroscience. Most recently, since October 2014, *Nature* journals require a statement whether (and where) the code underlying research results is available (“Code share” editorial, *Nature* 2014:514, p536). *Neuroinformatics* has a similar approach: authors can optionally include an information sharing statement noting whether resources such as code and data are available (Ascoli, 2014). Outside of the sciences, since its creation in 2005, the journal *Quarterly Journal of Political Science* has required all empirical papers to be reproducible.

In summary, the R computing environment provides many of the tools for reproducible research. Other languages, such as python, have equivalent notebook tools to help researchers share their work. This approach to writing papers is a powerful way of sharing one data and results with others. It allows others to easily verify the results, and then directly use the code and data associated with the article. This should lead us to a more cooperative environment for sharing our research.

## Acknowledgements

I thank Professors Joseph Corbo, Abbie Hughes, David Marshak and Heinz Wässle for providing the retinal mosaics included in this paper. Thanks to Ben Marwick and Sarah Mount for help with Docker testing and comments on the paper.

Session information
- R version 3.2.0 (2015–04–16), x86_64-apple-darwin13.4.0
- Locale: en_GB.UTF-8/en_GB.UTF-8/en_GB.UTF-8/C/en_GB.UTF-8/en_GB.UTF-8
- Base packages: base, datasets, graphics, grDevices, methods, stats, utils
- Other packages: knitr 1.11, sjedist 0.1, sjedmin 0.21, sjevor 0.32, sp 1.2–0, spatstat 1.42–2, splancs 2.01–37, xtable 1.7–4
- Loaded via a namespace (and not attached): abind 1.4–3, deldir 0.1–9, evaluate 0.7.2, formatR 1.2, goftest 1.0–3, grid 3.2.0, lattice 0.20–33, magrittr 1.5, Matrix 1.2–2, mgcv 1.8–7, nlme 3.1–122, polyclip 1.3–2, stringi 0.5–5, stringr 1.0.0, tensor 1.5, tools 3.2.0

